# Identifying progressive gene network perturbation from single-cell RNA-seq data

**DOI:** 10.1101/297275

**Authors:** Sumit Mukherjee, Alberto Carignano, Georg Seelig, Su-In Lee

## Abstract

Identifying the gene regulatory networks that control development or disease is one of the most important problems in biology. Here, we introduce a computational approach, called PIPER (<u>P</u>rogressIve network <u>PER</u>turbation), to identify the *perturbed genes* that drive differences in the gene regulatory network across different points in a biological progression. PIPER employs algorithms tailor-made for single cell RNA sequencing (scRNA-seq) data to jointly identify gene networks for multiple progressive conditions. It then performs differential network analysis along the identified gene networks to identify master regulators. We demonstrate that PIPER outperforms state-of-the-art alternative methods on simulated data and is able to predict known key regulators of differentiation on real scRNA-Seq datasets.

## I. Introduction

Recent advances in the field of single cell RNA sequencing (scRNA-seq) have enabled us to ask novel biological questions, which creates needs to develop new statistical methods to address them [1]–[3]. Unlike bulk RNA-seq or microarray measurements, scRNA-seq captures cell-to-cell variability in gene expression programs. This inter-cellular variation holds the key to inferring how genes transcriptionally regulate each other (i.e., gene regulatory network) and how their expressions and interactions change across cell states. Several studies have recently taken advantage of this data to examine biological processes such as differentiation [4]–[6] in a range of cell types and organisms. In this paper, we present the PIPER approach that aims to infer *progressive network changes* across different cellular states (e.g., differentiation) and the *regulator genes* whose connection with other genes are significantly different between the network estimates. The regulator genes are interpreted as *genes likely to have driven the network differences* (Figure 1).

**Fig. 1.**
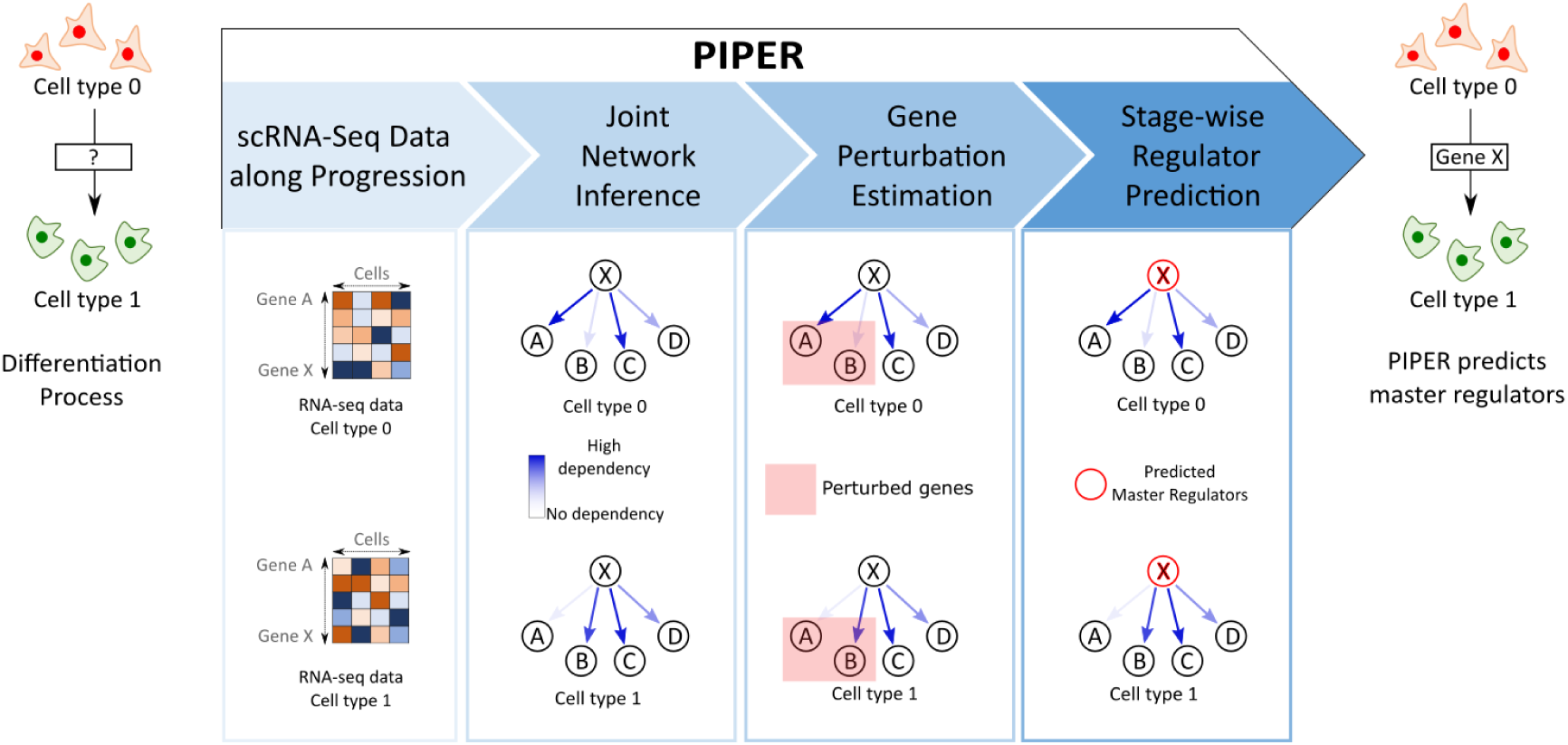
An overview of the different steps involved in master regulator prediction by PIPER. The input to PIPER is scRNA-Seq data from different states in a biological progression as seen in the cartoon on the left. The output of PIPER is the predicted set of key regulators for each stage of the progression as seen in the cartoon on the right.

PIPER has three unique advantages over existing approaches: First, although the aforementioned studies have been able to identify gene expression changes during differentiation, existing methods are not designed to identify *key regulators* of these processes and provide mechanistic explanations. Second, PIPER extends existing differential network analysis methods that have been successful at identifying key regulators of disease by detecting changes in gene network structures between different conditions [7]. Unfortunately, existing methods are intended to do pairwise comparisons between two independent conditions instead of capturing *structured* changes in multi-step or branched progressions. This limitation had not been an issue for bulk RNA-seq or microarray data, where high-volume and high-granularity data from multiple different conditions were rarely available. We need to extend differential network analysis methods to handle *multiple structured* conditions to leverage these data. Finally, PIPER uses the underlying distribution of scRNA-seq data that is more suited to count-valued sequencing data rather than normalized microarray data or bulk RNA-seq data.

PIPER extends previous works on network inference and differential network analysis method in the following ways. (1) *Network inference:* The earliest approaches for identifying gene networks computed correlations or mutual information between genes and preserved interactions only above a threshold value [8]. While simple, such methods often lead to the identification of spurious interactions caused by indirect interactions. This drawback is overcome in partial correlation networks [9]. A complimentary approach uses Gaussian graphical models (GGM) [10] to estimate the *conditional dependencies* between genes, assuming that the data follow a Gaussian distribution. Modifications of these methods make use of prior knowledge about the graph degree distribution [11], block structure [12] and pathway information [13] in order to further improve network inference. In practice, better performance has been achieved when data from multiple conditions are used to jointly estimate the networks at each individual condition [14], [15] instead of estimating them separately. To take the maximal advantage of scRNA-seq data, recent work [16] has focused on developing network estimation methods specifically suited for such data and has demonstrated their superior performance when compared to conventional methods on count-valued data. PIPER extends these approaches by using a local Poisson graphical model with a penalty that enforce consecutive states in a progression to have similar graph structures. PIPER thus provides a scalable algorithm for learning a genome-wide network from count-valued data. (2) *Differential network analysis:* The most direct approach to solve this problem is to identify genes whose connections change extensively between pairwise conditions. While efficient, this method relies heavily on the accuracy of the network structure learning process. DISCERN [17], a more general framework that utilizes the network parameters (edge weights) instead of network structure, was shown to be robust to errors in network structure estimates. PIPER extends DISCERN by identifying *both* perturbed genes and key regulators from *multiple structured conditions*. PIPER utilizes both perturbation analysis as well as jointly estimated network structures to identify highly connected genes which undergo strong perturbations between different conditions. Additionally, unlike DISCERN, these individual steps of PIPER are tailor-made for *scRNA-seq data* by using a Poisson distribution.

The main contribution of this paper is the development of a general statistical method that automatically predicts key drivers of progressive gene network changes, using as input scRNA-seq data measured in multiple points over the progression.

## II. METHODS

### A. Identifying conditional dependencies between genes

Count-valued scRNA-seq data have been successfully modeled by using the Poisson graphical models [16]. The paper proposed the use of a local model for each node, called the Local Poisson graphical model (LPGM), followed by combining information from each of these local models to infer the structure of the whole network. This is equivalent to solving the following optimization problem:

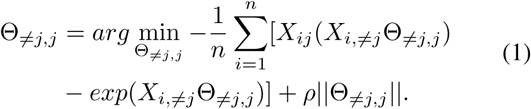

Here, the *j*th column of Θ ∈ ℝ^*g*×*g*^ contains the LPGM weights to model the expression level of gene *j* based on all the other genes. Specifically, Θ _≠*j,j*_ ∈ ℝ^(*g*−1)^ represents the *j*th column of Θ except the *j*th element. 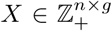 is a matrix of observed count values having *n* sample points and *g* genes, *X*_*ij*_ represents the value of the *i*th observation of *X*_*j*_, the subscript notation ≠ *j* represents a sub-matrix of the original matrix which contains all rows/columns except the *j*th one (depending on where it appears in the subscript). Finally, *ρ* ≥ 0 is the regularization parameter controlling the amount of sparsity in Θ.

To leverage the availability of data from multiple structured states, PIPER extends the estimation step of the LPGM formulation to multiple conditions as seen in equation 2.

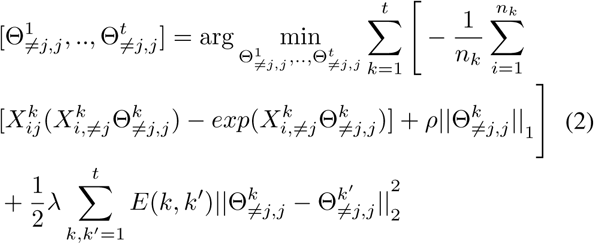

This formulation simply estimates the parameters for all conditions (Θ^1^…Θ^*t*^) together and has an additional regularization term *λ* which penalizes the 2-norm distance between networks estimated at neighboring states (*k* and *k*′). *E*(*k*, *k*′) is an indicator function whose value is 1 if the states *k* and *k*′ are adjacent, otherwise it is 0. We assume that the adjacency of the observed states is known to the user *apriori*. In the biological context, this modification allows us to model several types of progressions such as cell type lineage trees, differentiation events or spatial dependencies. This technique has been successfully used in the past for GGM’s [14] and is shown to result in fewer spurious network linkages.

### B. Identifying highly perturbed genes between networks from different states

Identification of genes whose dependencies change between two conditions has been done in the past by DISCERN [17], an algorithm that ranks nodes whose regulators have been perturbed the most. However, in its original formulation [17], DISCERN was not explicitly designed to work with scRNA-seq and hence made an assumption of a Gaussian distribution for the data. Here we have adapted this method for count-valued processes by assuming that the data has a Poisson distribution.

Scoring perturbed genes requires first identifying the gene regulatory network for two separate conditions followed by calculating the perturbation score. The perturbation score captures how well conditional dependencies of genes in one condition can explain the data for the other condition. The perturbation score is defined as follows:

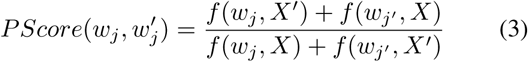

Where *w*_*j*_ and *w*_*j*′_ are the vectors of regulator weights for gene *j* in conditions 1 and 2. While *X* and *X*′ are the data sets for conditions 1 and 2 respectively. *f* (*w*_*j*_, *X*) is the unpenalized local Poisson graphical model cost function as seen in equation 1 (with *ρ* set to zero).

### C. Identifying clusters of similarly perturbed genes

While there exist methods to identify perturbed nodes between two conditions, they cannot be directly extended to multiple conditions particularly in the form of a biological progression. Here we propose identifying the clusters of similarly perturbed genes in order to study genes potentially regulating them.

In order to identify clusters of similarly perturbed genes, we first normalize the PScore of each gene across all conditions to have a 2-norm of 1. The normalized score for each gene is then tiled at all conditions to estimate the relative perturbation between conditions. The normalization for each gene also ensures that all scores have roughly the same scale, which is important for clustering and, moreover, it ensures that condition-specific scores for genes that are predicted to be highly perturbed at all conditions are low. Such genes are unlikely to be master regulators since their perturbations are expected to be condition specific. Next, we may perform an optional step of setting a threshold value for the normalized PScore to separate out genes that are not perturbed at any condition. This step is not seen to be important for small networks but is crucial for larger networks. Finally, we apply k-means clustering to obtain clusters of genes that have similar perturbation patterns across different conditions. The appropriate number of clusters can then selected by using well known methods such as the ‘elbow method’ [18].

### D. Sparsifying network to improve specificity

Our network estimation method leads to the estimation of several networks at each sparsity level (corresponding to each value of *ρ*). For the purposes of this paper, we have estimated several networks in the *ρ* = [0.01, 10] range. The upper and lower cut-offs for *ρ* were empirically selected by observing the structure of estimated networks. In order to only retain edges we are confident about, we identify the most conserved network structure between the networks estimated at different values of *ρ*. This is done by first constructing an unweighted adjacency matrix for each value of *ρ* (*A*_*ρ*_(*i, j*) = *I*(|Θ_*ρ*_(*i, j*)| > 0), where *I*(.) is the indicator function) and then taking the elementwise product of the the adjacency matrices estimated for all values *ρ* as follows:

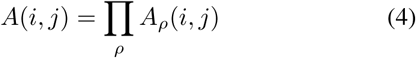

This network is then further sparsified by eliminating edges of which the values of correlation between the two genes is lower than a threshold value. For the purposes of this paper, we empirically set this threshold to be the mean of the correlations between all genes in the network. While sparsifying the network might lead to the elimination of some actual conditional dependencies, we argue that specificity is more important to our problem of master regulator identification than sensitivity. It must also be noted that unlike in [16], we do not force *A* to be symmetric, thereby allowing it to represent a directed graph.

### E. >Identifying key regulators with temporally or pseudotemporally hierarchical data

When the different conditions of the data set are in the form of a temporal/pseudotemporal hierarchy, it roughly corresponds to a process where we have access to intermediate stages of differentiation or development. Such a process can be represented by a directed graph. Each state *i* will also have an associated sparsified incidence matrix *A*^*i*^. In such a case, for each state node in the hierarchy, *i*, we first find the highest valued PScore cluster(s) relevant to this state and collect the indices of the genes in these cluster(s) into a set *S*. We then find the number of times each gene in our data set is predicted to be a regulator of the genes (in the set *S*) in the state *i* as:

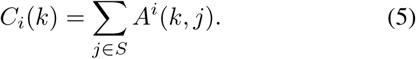

Here, *C*_*i*_(*k*) is the number of highly perturbed genes targeted by gene *k* in state *i*.

### F. Identifying key regulators from snapshot data with missing intermediate states

For some of the scRNA-seq data sets, the data are available from a single time point containing only different mature cell types. In such a case, to identify the key regulators in the progenitor cell states that lead to each of the different mature types, one has to infer them from the observed cell types. A natural way to do this could be to look at the observed daughter cell types of a unobserved progenitor cell type and identify the highly perturbed hub genes between the different conditions. Here we propose a strategy to do this when the structure of the differentiation tree is known. First, we construct a graph joining the closest observed cell state where distance is measured by number of nodes along the tree separating a pair of nodes (we assume the true structure is known apriori). The perturbation analysis is then performed for each pair of neighbors (*i, j*) for this new graph. We then find the cluster for which the mean PScore at the pairwise condition (*i, j*) is highest and consolidate the genes in the cluster into a set *S*. Lastly we find the number of times each gene in our data set are predicted to be regulators of the genes in pairwise condition (*i, j*) as:

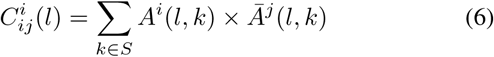

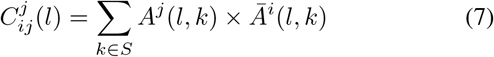

Where 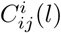 is the number of highly perturbed genes targeted by gene *l* in pairwise-condition (*i, j*) that are specific to state *i*. *A*^*i*^ is the incidence matrix at state *i* and 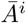 is the complement of the incidence matrix.

## III. RESULTS

### A. Simulation study I: Testing the efficiency of perturbed node estimation

To test the accuracy of the perturbed node estimation process, we first generated synthetic Poisson distributed data from random graphs with 30 nodes and 100 samples using the methodology described in [16]. To simulate different biological conditions, we randomly deleted 5% of the edges from the original graph. Graphs of different sparsity levels were generated by varying the connection probability between pairs of nodes. 20 random graphs were generated at each sparsity level. We also used these data to demonstrate the usefulness of joint network estimation by testing against a variant of PIPER where the network similarity penalty *λ* was set to 0. The ranking of perturbed genes for PIPER is obtained by sorting genes according to their PScore calculated between the two conditions. The accuracy at each sparsity level is measured by calculating the mean of the fraction of the *N* perturbed genes that are ranked in the top *N* genes according to the different methods over all trials (and values of *ρ* for PIPER and Treegl). We then compare PIPER against various other specialized methods for identifying perturbation in networks namely DISCERN, Treegl (with perturbation identified with DISCERN), D1 Score, LNS Score and ANOVA (the latter three are computed as described in [17]). For both PIPER and Treegl, the tuning parameter *λ* (that enforces similarity between successive conditions) is chosen to be the best performing one at each sparsity level. It can be seen in Figure 2a that for graphs of most sparsity levels, both formulations of PIPER outperform all competing methods while the joint network inference seems to have a higher accuracy than *λ* = 0 particularly for sparser graphs.

**Fig. 2.**
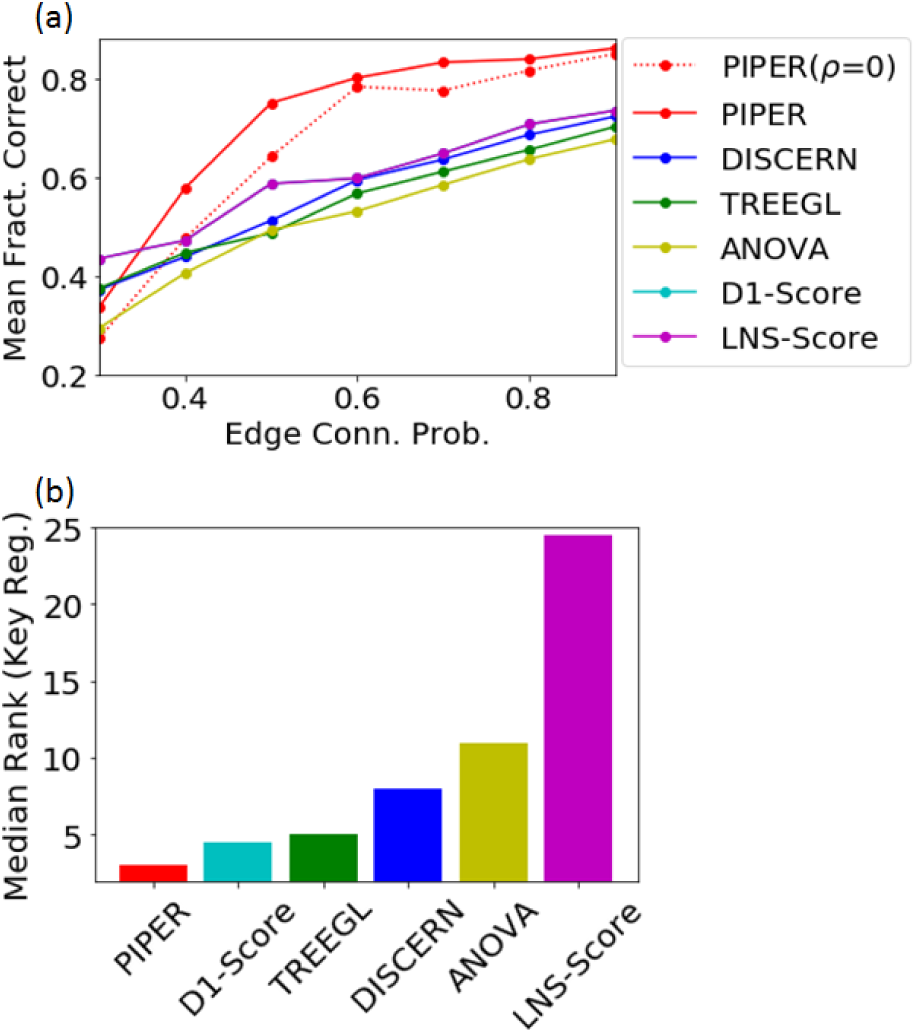
a) Testing accuracy of PIPER at identifying perturbed genes for graphs having different sparsity levels. The mean fraction of perturbed genes that are correctly predicted is calculated by averaging across all values of the sparsity parameter *ρ*. b) Comparison of predicted mean rank of actual key regulator for various methods on scale free graphs of degree 3. PIPER outperforms other methods in predicting the key regulator genes.

### B. Simulation study II: Testing key regulator identification on synthetic data from scale-free networks

Having demonstrated PIPER’s accuracy at identifying perturbed genes between networks from two different conditions, we test its accuracy at identifying the key regulators of a simulated biological progression. This is a three state system where a single progenitor state differentiates into two daughter states. The procedure for generating the graph, perturbed graph and the data sets is similar to the previous sub-section with the notable exception being that here we assume the graph is scale-free [19] (average degree = 3) i.e. its node degree distribution follows a power law. Scale–free networks are common in biology and have many nodes with small node degrees with a few nodes (called hubs) having much higher node degrees. It has also been demonstrated that most key regulators are hubs in their respective networks [20]. Hence, when deleting the edges, we first perform a weighted sampling (where the weights are proportional to the degree of the node) and then select one of the outgoing edges of the selected node, uniformly at random to delete. This method ensures that hub nodes are more likely to be the main regulators for the perturbed genes. Due to the randomness of the edge deletion, the two daughter networks are also different from one another. We now use this method to generate 20 random set of networks (and data sets) of the same dimension as the previous subsection with 10% of the edges deleted in the perturbed network. We then proceeded to first obtain the PScores for all genes, and the regulators of the genes with the top *N* PScores, where *N* is the number of actually perturbed genes (as in the previous sub-section). We then ranked the predicted regulators of these genes according to their frequency of occurrence. For each trial, we then used this ranked list to find the predicted position of the actual top regulator for each branch of the differentiation process and calculated the average median rank across both branches. We found that PIPER outperformed other algorithms in identifying the key regulators when compared by the median predicted rank of the actual top regulator (as seen in Figure 2b).

### C. Case I: PIPER correctly identifies early and late regulators of mouse embyonic stem-cell (mESC) differentiation

For a first application we used scRNA-seq data [1] collected at four time points (day 0, 2, 4, 7) during differentiation of mouse embryonic stem cells to epiblast cells. We assume data from each day represents a different dominant cell state during the differentiation process. We focused on a panel of 89 genes comprising of essential housekeeping genes, key regulators of mouse ES differentiation and differentiation markers [21]. As per the PIPER workflow, we first performed a joint network inference, followed by gene perturbation estimation across successive conditions and clustering of genes on the basic of PScores. This process lead to the identification four distinct clusters of genes (Figure 3b).

**Fig. 3.**
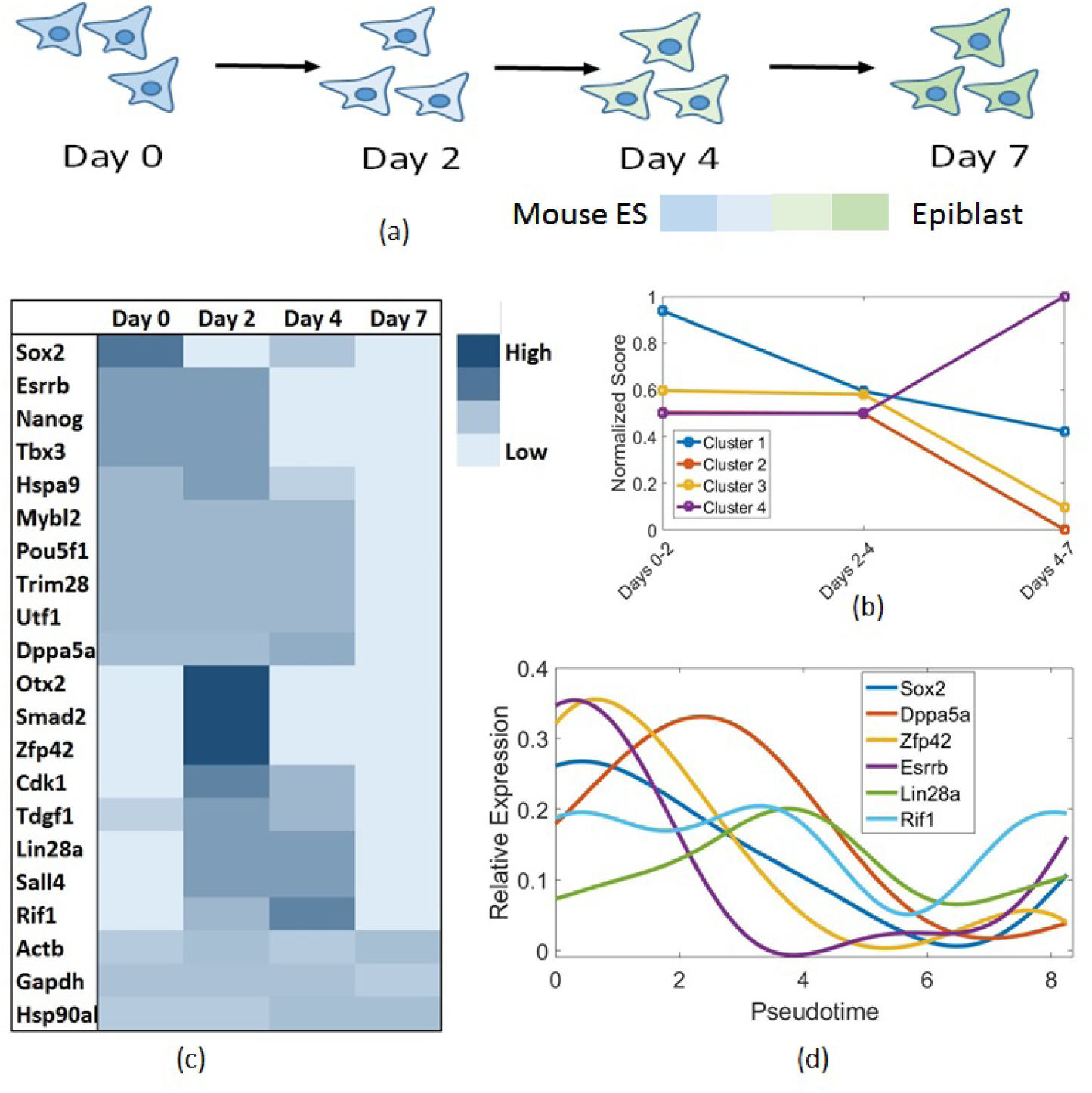
time points from which the data is available. b) Clusters of P-scores across multiple days. c) Heatmap of number of different perturbed genes targeted by the predicted key regulators of differentiation across different days. d) Expressions of different predicted key regulators plotted against pseudotime inferred from Monocle. Peaks in expression patterns are closely match the stages on which the regulators are predicted to act.

Next we identified the clusters that are relevant to each day (by looking at the cluster that contained the highest PScore for a particular day): cluster 1 for day 0, clusters 1 and 3 for day 2, clusters 3 and 4 for day 4, and cluster 4 for day 7. For each day we now only look at the genes from the clusters relevant to the day and identify their regulators (shown in Figure 3c). The column normalized heat map shows how many perturbed genes are targeted by each regulator across different days.

We note that most predicted regulators from early stages (days 0, 2 and 4) are known differentiation regulators. For example, Pou5f1, Sall4, Zfp42, Utf1, Sox2, Nanog, Esrrb, Dppa5a, Rif1 are all known to have roles in maintaining pluripotency [22]–[26]. Of these genes, Nanog and Sox2 have been widely reported to be among the earliest regulators of mouse ES differentiation [26], [27] and have been correctly implicated by PIPER to control early stages of differentiation (days 0 and 2, respectively). Moreover, Rif1, which is predicted to regulate subsequent stages of differentiation (days 2 and 4) is known to be activated by Nanog [26], consistent with PIPER’s prediction. Since most of the differentiation process is completed by day 4, we note that the predicted regulators from day 7 are all housekeeping genes.

To further validate the accuracy of the order of regulator action, we used Monocle [4] to order cells by differentiation progress. This cell state information can then be used to plot gene expression as a function of pseudotime (an arbitrary metric representing progression along cell states). While Monocle is not designed to identify the key regulators of differentiation, one would expect the expression patterns of our predicted regulators for different stages of differentiation to show big changes during the predicted period of their effect. This can be seen in Figure 3d, hence providing a further validation to PIPER’s predictions.

### D. Case II: PIPER identifies genes responsible for lineage formation in neuronal cell types

To test the accuracy of PIPER on branched snapshot data, we looked at a data set [2] containing neurons and non-neuronal cell types from the mouse cortex and hippocampus. We focused on a sub-set of the dataset containing 3 mature cell types deriving from a common precursor (namely astrocytes, oligodendrocytes and interneurons, Figure 4a). We aim to identify the key regulators that drive differentiation of the common neuronal stem cell precursor into each of the three terminal cell types.

**Fig. 4.**
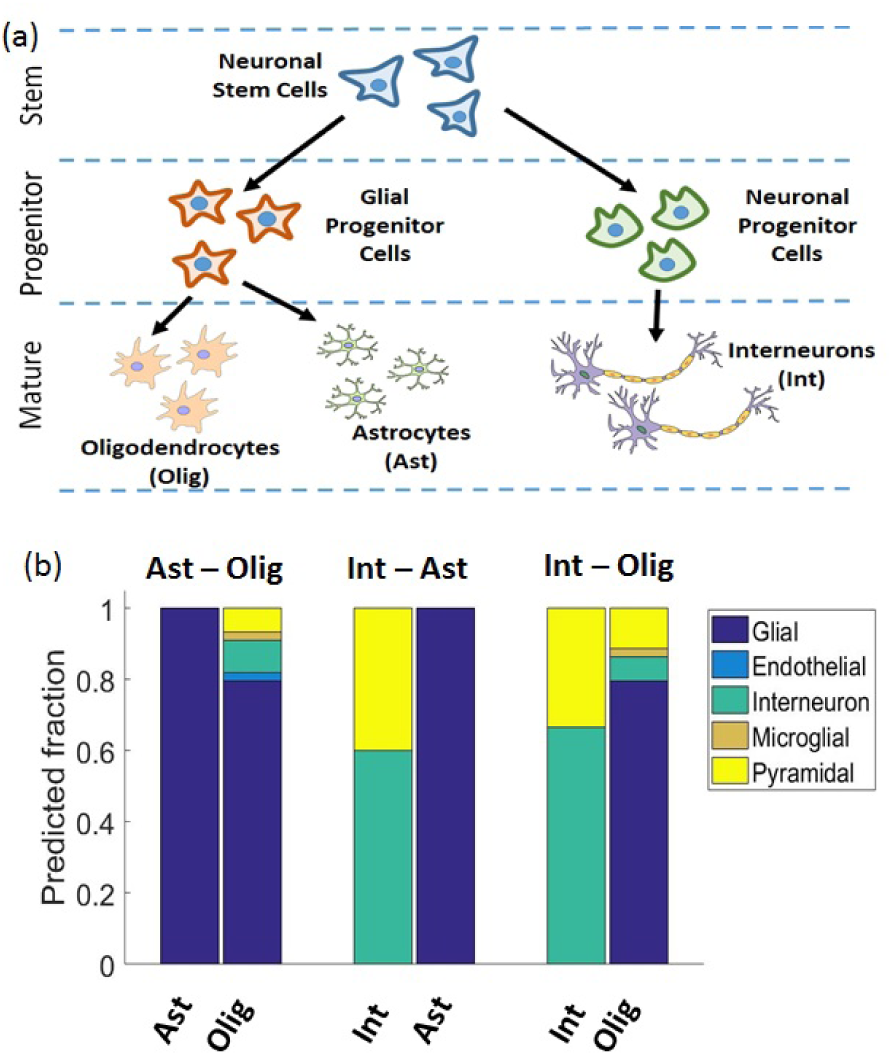
Application of PIPER to scRNA-seq data obtained from mouse brain samples correctly identifies genes enriched for different neuronal cell types. a) Neuronal stem cell lineage tree. b) Fraction of predicted regulators of each stage belonging to different progenitor cell types. It can be seen that the regulators of most daugther cell types are specific to the correct progenitor type.

We first identified a subset of genes that are of interest for differentiation. We started with a list of over 3,000 genes that were reported in [2] to be uniquely upregulated in any one of the main cell types. We then limited the list to 528 by excluding genes that do not have at least one count on average in one of the cell types. This filtering step removed genes that have too few counts to lead to a meaningful identification of network structure. We then performed network estimation with all three cell types and estimated the perturbation of each cell type compared to their closest neighboring cell type along the differentiation tree. Upon normalization of Pscores and clustering, we were able to obtain three major clusters of genes which are upregulated in at least one of the cell type pairs. After thresholding, we were left with only 68 out of 528 original genes. Next, we identified the regulators of these genes by sparsifying the networks at each condition as previously described in the Methods section. We thus obtained the list of key regulators by retaining genes which have regulate more than one perturbed gene. To check the validity of our results we compared against the list of genes unique to each cell type obtained from [2]. The genes specific to each progenitor type is obtained by merging the list of highly expressed genes for each of their daughter types. We note that while this is an approximate way to obtain genes specific to progenitor types, a more comprehensive list would require data from the progenitor states themselves. Figure 4b shows the fraction of predicted regulators unique to each of the two conditions that overlap with genes associated with different progenitor types according to [2]. It is interesting to note that not only are most predicted regulators also associated with correct progenitor cell types, when they do overlap with other cell types it is usually with a closely related cell type. Moreover, several of PIPER’s predicted key regulator genes such as Dbi, Gsn, Olig1, Lgi3 or Tcf4 are known to be important differentiation regulators of neuronal cell types. Several of the genes identified by PIPER such as Atp2a2, Mal, Hsd17b7, or Pdlim2 are known to play roles in differentiation in other cell types. These genes could be subject of future experimental study to determine their role in neuronal differentiation events.

## IV. CONCLUSIONS

In this paper we presented PIPER, a tool for identifying key regulators of differentiation events from scRNA-seq data. We demonstrated that PIPER can deal with various forms of data, such as time-series data containing intermediate states as well as data from one time point representing mature differentiated cell types. We benchmarked the individual components of PIPER on synthetic data and tested the complete workflow on two different biological data sets. PIPER predicted known key regulators of each differentiation/development process and notably predicted the correct temporal ordering of regulator action in mouse ES differentiation.

Future work can be aimed at inferring the graph structure of differentiation states along with the prediction of key regulators. This change would address a current limitation of PIPER, which is the need for an accurate differentiation graph to be provided to the algorithm. While this graph might be easy to obtain for well-studied processes, a more general approach could enable the study of new processes about which enough information is not available.

